# Effects of the Mu Opioid Receptor Positive Allosteric Modulator BMS-986122 On Opioid Efficacy in Rat Neuropathic Pain States

**DOI:** 10.64898/2026.05.03.722511

**Authors:** Benjamin M. Clements, Ipek Berberoglu, Katherine L. Burke, Stephen W.P. Kemp, John R. Traynor

**Author notes:** Corresponding author., Department of Pharmacology, University of Michigan Medical School, 220C MSRB III.1150 West medical center Drive, Ann Arbor, MI 48109-5632.

## Abstract

**Background:** Neuropathic pain is a major source of disability and distress with few pharmacological options for treatment. Opioid drugs can be effective, but high doses are needed, leading to unwanted effects. BMS-986122 is a positive allosteric modulator of the mu opioid receptor that potentiates acute opioid antinociception without increasing opioid-induced constipation, reward, or respiratory depression. Therefore, we asked if BMS-986122 could increase the effects of low-dose opioid analgesics in chronic neuropathic pain.

**Methods:** We employed the spared nerve injury and tibial neuroma models in rats and assessed the tactile hypersensitivity of the hind paw and site of neuroma, respectively.

**Results:** Administration of low doses of (R)-methadone, morphine, or buprenorphine slightly reduced the tactile hypersensitivity of the hind paw the in spared nerve injury model. Pretreatment with BMS-986122 significantly enhanced the reversal of hypersensitivity, reaching the effect of high-dose gabapentin, a standard of care in neuropathic pain. Pretreatment with BMS-986122 similarly increased the anti-allodynic effects of low dose (R)-methadone on neuroma pain. A similar effect of (R)-methadone in the absence of BMS-986122 was only observed at a dose where respiratory distress was seen.

**Conclusions:** These findings show that allosteric modulators of the mu opioid receptor such as BMS-986122 can enhance opioid activity that could translate to a safe and effective treatment for chronic neuropathic pain.

## Introduction

Chronic pain remains severely undertreated, affecting an estimated 30% of people worldwide.^1^ The opioid drug class represents some of the most powerful pharmacological tools to treat severe pain, but their widespread use is limited by tolerance, gastrointestinal effects, abuse liability, and respiratory depression.^2^ Chronic neuropathic pain affects 7-10% of the population^3^ and is considered resistant to opioid therapy, requiring high opioid dosing in patients with resultant unwanted effects.^4^ This limits pharmacotherapy options and contributes to the undertreatment of patients with neuropathic pain. This is especially true since first-line treatments like gabapentin and tricyclic antidepressants show efficacy only in a subset of patients.^5,6^

Clinically useful opioid drugs exert their actions by binding to the orthosteric site on the mu opioid receptor (MOR). Recently, positive allosteric modulators (PAMs) of MOR have been suggested as a way to provide pain relief either as standalone treatments or as adjuncts to traditional opioids providing an opioid sparing effect.^7–9^ PAMs interact with the receptors at a location separate from the orthosteric site to increase the potency and/or efficacy of opioid agonists. PAMs for MOR, such as BMS-986122, have been shown to enhance the *in vitro* activity of both opioid drugs and endogenous opioid peptides.^10^ In rodent models of acute antinociception, BMS-986122 increases the potency of clinically used opioids and endogenous opioid peptides, without enhancing opioid-induced constipation, reward, or respiratory depression.^7,11^ These preclinical data suggest that use of allosteric modulators may also alter the activity of opioids in a chronic pain state with a substantial increase in the therapeutic index, making opioids more efficacious and safer for chronic neuropathic pain patients.

Several rodent models of nerve injury have been used to investigate the efficacy of analgesic drug treatment. The spared nerve injury model (SNI) has been widely adopted as a model of mononeuropathy. This injury results in non-resolving pain behaviors that closely mimic clinical phenotypes of neuropathic pain, particularly hypersensitivity to tactile stimuli.^12^ This hypersensitivity, in both patients and SNI animals, can be reversed with gabapentinoids, serotonin-norepinephrine reuptake inhibitors, and opioids, albeit at a reduced potency compared to their action in acute pain states.^13,14^ However, following nerve injury, attempts at nerve regrowth occur and can lead to the formation of a neuroma, a large mass of unorganized axon fibers and non-neuronal tissues.^15^ The neuroma, when symptomatic, is extremely sensitive to tactile stimulation and cold, and can elicit spontaneous pain responses.^16^ Symptomatic neuromas are common following amputation, with occurrences ranging from 40% to 70%.^17,18^ Such neuromas are generally treated through surgical methods,^19^ but in conditions where surgery is not appropriate, pharmacotherapies must be employed, although these have been largely limited in effectiveness, including first-line treatments like gabapentin and amitriptyline.^20,21^ Morphine has shown limited success when administered intravenously,^22^ but assessment of high-dose oral morphine showed no difference from placebo.^23^ Improvements in pharmacological strategies for this patient population could provide meaningful benefits, as well as increase treatment options for additional neuropathic pain states, as neuromas are an expected result of any nerve injury.^15^

To test the efficacy of opioid PAMs in chronic neuropathic pain, BMS-986122 was used as a pretreatment to the MOR agonists (R)-methadone, the active enantiomer of racemic methadone, and morphine following SNI treatment in rats. BMS-986122 was selected as it has been well characterized *in vitro*^10,24,25^ and *in vivo*,^7,11^ and enhances opioid-mediated antihypersensitive effects in inflammatory pain models, suggesting it could be effective in neuropathic pain. We also studied the partial agonist buprenorphine. Due to the risks associated with full agonists, the use of partial opioid agonists is becoming more preferable in the clinic.^26^ In the neuroma model BMS-986122 was also used as a pretreatment to (R)-methadone. We chose methadone for these experiments as it is the most sensitive opioid to PAM activity in *in vitro* ^27^ and *in vivo* in acute pain models.^11^

Results show the reversal of nerve injury-induced tactile hypersensitivity by the opioids, including buprenorphine, in either model was greater in the presence of BMS-986122, allowing sub-effective, and therefore safer, doses of opioid to be used. This indicates opioid PAMs could be used to increase opioid efficacy in various forms of neuropathic pain and provide additional therapeutic options for patients.

## Materials and Methods

### Experimental Design

Behavioral assays were performed using a repeat measures model to better accommodate animal welfare and large numbers of animals undergoing unnecessary surgeries and experiencing neuropathic pain states. Doses were applied in an escalating manner, with 48 hours of washout in between assays to prevent tolerance and residual drug effects. At the end of a treatment group, vehicle tests were performed to examine order effects or learned behaviors. In addition, a replicate study was performed at the end of testing to confirm behavioral effects. By employing a repeat measures model, groups of 8 animals were appropriate for all tests. Equal numbers of males and females were used for all studies.

### Drugs and Reagents

R-methadone and buprenorphine were generously provided by the NIDA Drug Supply (Bethesda, MD USA). Morphine sulfate was purchased from Spectrum Chemical (New Brunswick, NJ USA). Gabapentin was purchased from Sigma-Aldrich (St. Louis, MO USA). BMS-986122 (BMS-122) at > 96% purity was synthesized at the Vahlteich Medicinal Chemistry Core at the University of Michigan as previously described.^10^ Opioid agonists were dissolved in 0.9% sterile saline to the appropriate concentration to provide a 1 mL/kg dose. BMS-986122 was dissolved in DMSO and constituted in a vehicle comprised of 10% DMSO, 10% ethoxylated castor oil, and 80% sterile H_2_O. All drugs were administered subcutaneously.

### Animals

For all behavioral studies, male and female Sprague-Dawley rats were used. All rats were two months of age at the beginning of surgery. Rats were group housed in accordance with sex, maintained on a 12:12 hour light/dark cycle, and allowed *ad libitum* access to food and water. Experiments were approved by the University of Michigan Committee on the Use and Care of Animals. Experimental procedures described in this study were conducted in strict accordance with the guidelines set forth by the Unit for Laboratory Animal Medicine (ULAM) at the University of Michigan and adhered to the National Institutes of Health (NIH) Guide for the Care and Use of Laboratory Animals to ensure ethical standards, minimize animal use, and reduce discomfort.

All behavioral tests were performed with repeated measures, with assessments before induction of the pain state and after multiple doses within the same animal. As such, animals were tested, maintained, and monitored closely for a minimum of three weeks following the start of behavioral assays. New baseline measurements were determined prior to each drug treatment and compared to check for potential changes in normal tactile sensation that might have been caused by repeat treatments. The structure of repeat measures paradigm was designed to limit order effects, and a minimum of 48 hours washout was allowed between behavioral tests.

### Surgical Procedures for Injury Models

#### Induction of Tactile Hypersensitivity via Spared Nerve Injury

Spared nerve injury was performed as described in Decosterd and Woolf^12^ with minor variations. Eyes were treated with ophthalmic ointment to prevent dryness, and aseptic surgical techniques were utilized throughout the procedure. The mouth was cleared of any obstructions, and body temperature was maintained using a heating pad to ensure normothermia. Pain medication was provided in the form of carprofen at a dose of 4 mg/kg subcutaneously. Surgery was performed on the left thigh, which was marked accordingly. The lateral surface of the left thigh was shaved with a razor blade and cleaned alternately with povidone-iodine prep pads and alcohol wipes. Inhalation anesthesia was induced and maintained with 5% and 2% isoflurane/oxygen, respectively, and the rat was positioned in a left lateral decubitus position. A single, small linear incision approximately 3–4 cm long was made at the mid-thigh level along the lateral thigh, angled 15–20 degrees relative to the femur. Blunt dissection was performed through the biceps femoris muscle to expose the sciatic nerve and its three branches: the sural, common peroneal (CP), and tibial nerves. The tibial and CP nerves were carefully ligated using 6-0 vicryl sutures, 2 mm distal to the bifurcation, and approximately 3 mm of the distal stumps were excised (neurectomy) distal to the ligation point to prevent nerve regrowth. The sural nerve was preserved, avoiding any unnecessary contact or stretching. The incision was closed layer by layer using muscle and skin sutures. Throughout the procedure, minimal retraction was applied to reduce nerve stress, and bleeding was controlled with pressure as necessary.

Postoperative pain medication was administered for the first two days (4 mg/kg carprofen, s.c.). Postoperative check-ups were conducted to monitor any abnormalities, and sutures were removed after 10 days. Behavioral testing began in the maintenance phase of the pain state, 14 days after surgery.

#### Induction of Tactile Hypersensitivity via Neuroma Formation

Surgeries were performed on the right hind limb using aseptic techniques in accordance with regulatory protocols as described above. The tibial nerve was selected for creating neuromas due to its easy accessibility and isolation, as described previously.^28^ Rats were anesthetized using 2% isoflurane/oxygen and placed on a heating pad. The skin incision was made on the lateral aspect of the thigh followed by blunt dissection of the biceps femoris muscle. The tibial nerve was identified, isolated, and subsequently cut distally directly before entry into the posterior compartment of the leg. The proximal segment was relocated over the biceps femoris muscle, and the epineurium was sutured to the overlying dermis. The overlying skin was then tagged with a non-absorbable suture and marked with a pen to be used as a clear target for direct neuroma stimulation during behavioral testing. Behavioral testing began 8 weeks after surgery.

### Behavioral Tests

#### Tactile Sensitivity of the Hind Paw

Prior to SNI surgery, baseline tactile sensitivity of both hind paws was determined. Using various manual von Frey filaments, the 50% withdrawal threshold for both paws was determined using the Up-Down method.^29^ Fourteen days after injury, both the ipsilateral (left) and contralateral paw (right) were tested for tactile sensitivity. Following determination of the baseline, animals were administered BMS-986122 or vehicle. Fifteen minutes later, tactile threshold was again determined, and opioid agonist or vehicle was delivered. Following delivery of opioid agonist, tactile thresholds of both paws were determined at 15, 30, 45, 60, 90, and 120 minutes. Increasing doses and other treatment conditions were performed in the same animals, with a minimum of 48 hours to allow complete washout of any drug. Eight animals (4 males and 4 females) were used in each study.

#### Neuroma Score

Eight weeks after induction, the tenderness of the neuroma was determined using a variation of the clinically used Tinel test as described previously.^30^ In brief, one von Frey filament (15 g) was used to stimulate the skin above the neuroma (identified via mechanical assessment). The stimulation involved 5 gentle taps on the location of the neuroma. Stimulation ended either after the final tap (score of 6) or after a pain-like behavior was documented (fast withdrawal, flicking, and/or licking). This score is referred to as the “neuroma score”. A neuroma score was obtained prior to the start of any drug treatment, after BMS-986122 administration, and after agonist administration as described for the assays using the SNI animals. Repeat testing was also performed as described for the SNI animal experiments. Eight animals (4 males and 4 females) were used for all (R)-methadone studies. A total of 4 animals (2 males and 2 females) was used for the naloxone study.

### Statistical Analysis

Behavioral assay results were analyzed using Graph Pad Prism 10.2.2 (Boston, MA USA). For tactile sensitivity assays, agonist-vehicle, agonist-modulator, and vehicle-modulator groups were compared using a repeat-measure two-way ANOVA with Tukey’s multiple comparison test to examine for differences at each time point. These data are presented at mean ± S.E.M. No statistical power calculation was performed prior to the start of the study, and group sizes were determined based on experience with the models tested. For all assays, P < 0.05 was considered statistically significant. Animals were tested in sequential order. No animals were excluded from this study. The data were monitored for statistical outliers, but no data were excluded from the analyses presented. A total of 40 rats were used for this study.

## Results

### Spared Nerve Injury Induced Hypersensitivity

Patients with chronic neuropathic pain are known to have a reduced response to the pain-relieving effects of opioids.^4^ This reduction in analgesic efficacy is consistently seen in preclinical pain models.^31^ Based on its ability to increase opioid receptor activation *in vitro*^10^ and acute opioid-induced antinociception *in vivo*,^7^ the MOR-PAM BMS-986122 was assessed for its potential to enhance the effects of low dose opioids in the SNI model of mononeuropathy in the rat. A dose of 10 mg/kg of BMS-986122 subcutaneously (s.c.) was employed for all the studies since this is the highest dose possible due to solubility limitations of the molecule. All opioids and gabapentin were also administered via s.c. injection.

SNI greatly reduced tactile threshold in the hind paw measured by von Frey filaments. The 50% withdrawal threshold before surgery was 8.9 g and after surgery a hypersensitive response of 0.78 g was observed, as expected.^12^ Administration of 0.1 mg/kg or 0.32 mg/kg (R)-methadone did not alter the 50% withdrawal threshold. BMS-986122 alone also had no effect. However, pretreatment with BMS-98122 prior to (R)-methadone caused the opioid to show a dose-dependent decrease in tactile hypersensitivity (**Fig. 1A**, **1B**). A higher (R)-methadone dose of 1 mg/kg was able to fully reverse the tactile hypersensitivity, and this effect was not further enhanced in the presence of BMS-986122. (**Fig. 1C**). After SNI, the contralateral, uninjured paw responded to von Frey filaments with a 50% withdrawal threshold of 13.3 g (**Fig. 1D-F**). This increase in withdrawal threshold compared to before injury may be related to weight displacement, as the animal begins to limit pressure on the injured ipsilateral hind paw. (R)-methadone at 0.1 mg/kg and 0.32 mg/kg did not alter this threshold but 1 mg/kg (R)-methadone raised the threshold to 40 g. BMS-986122 did not affect the responses to methadone in this paw. (**Fig. 1D-F**).

**Figure 1.**
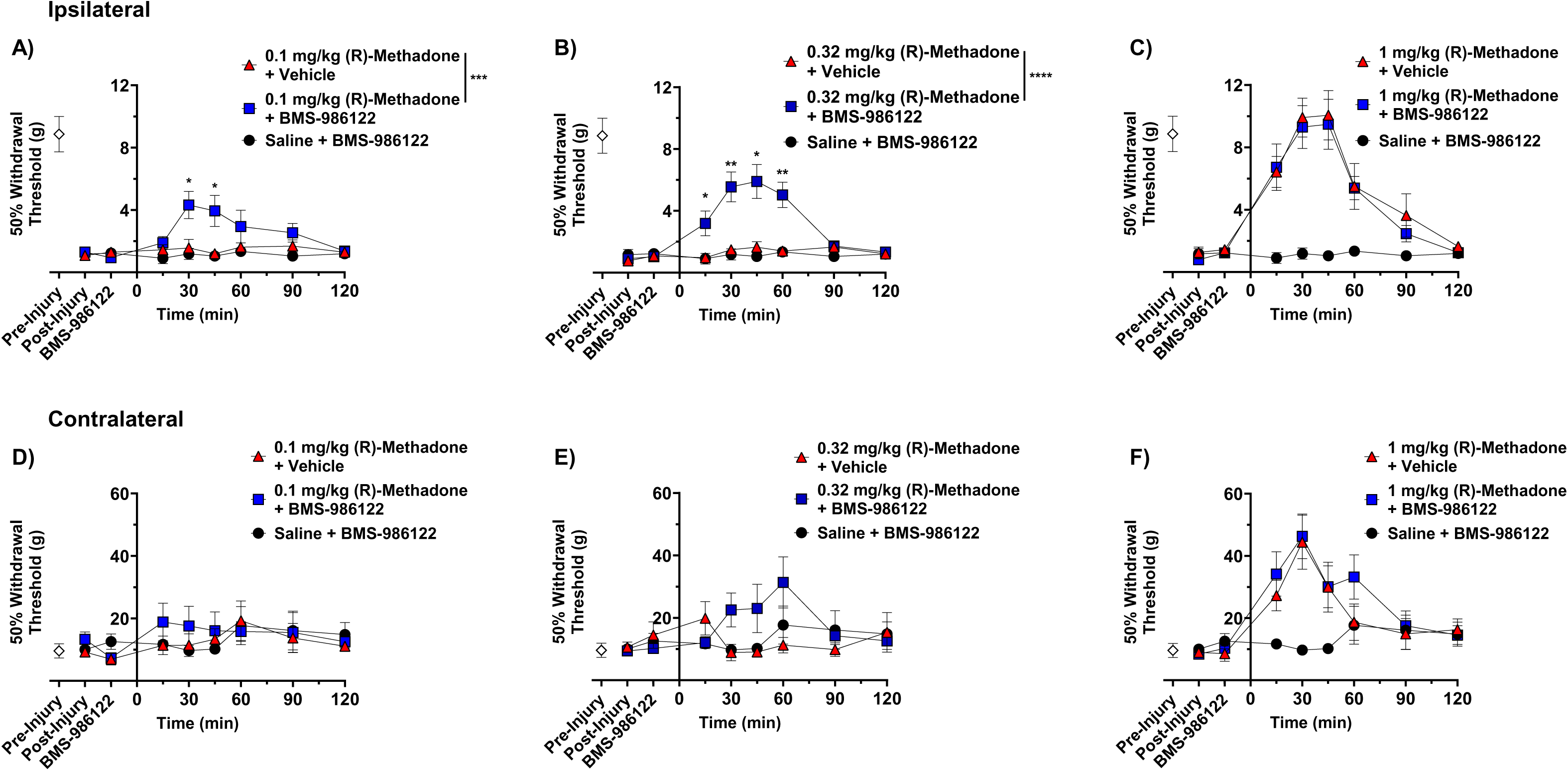
Effect of BMS-986122 on (R)-methadone-induced reduction in tactile hypersensitivity after nerve injury. Rats with tactile hypersensitivity in the injured hind paw after SNI received BMS-986122 (10 mg/kg, s.c.) as a 15-minute pretreatment to subcutaneous (R)-methadone at a dose of 0.1 mg/kg (**A**), 0.32 mg/kg (**B**), or 1 mg/kg (**C**). 50% Withdrawal thresholds were determined using von Frey filaments and the up-down method prior to injury (Pre-Injury) and prior to BMS-986122 administration (Post-Injury), 15 minutes after BMS-986122 administration (BMS-986122), and for 2 hours after (R)-methadone administration. Thresholds were simultaneously measured in the contralateral, uninjured hind paw after mice received 0.1 mg/kg (**D**), 0.32 mg/kg (**E**), or 1 mg/kg (**F**) (R)-methadone. Groups were compared using a repeat-measures two-way ANOVA with Tukey’s post hoc test, with significance reported between (R)-methadone in the presence of vehicle and (R)-methadone in the presence of BMS-986122 at each time point. * p < 0.05, ** p < 0.01, *** p < 0.001, **** p < 0.0001 (n = 8, (4 females and 4 males). Data are presented as mean ± S.E.M.

(R)-methadone, while a potent analgesic, is rarely used for this purpose in the clinic.^32^ Therefore, we tested whether the action of morphine could be enhanced by BMS-986122. Administration of low dose morphine (1 or 3.2 mg/kg) did not increase the 50% withdrawal threshold in the injured paw (**Fig. 2A, 2B**). At 10 mg/kg morphine showed a significant effect but did not fully reverse the hypersensitivity, indicating it was less effective than methadone (**Fig. 2C**). In contrast, the ineffective doses of 1 and 3.2 mg /kg morphine showed a significant antihypersensitivity action in the presence of 10 mg/kg BMS-986122 (**Fig. 2A, 2B**). The response to the higher dose of morphine was somewhat greater but this did not reach significance. Morphine showed an increase in tactile thresholds in the contralateral, uninjured paw, but this was not enhanced by BMS-986122 at any dose tested (**Fig. 2D-F**).

**Figure 2.**
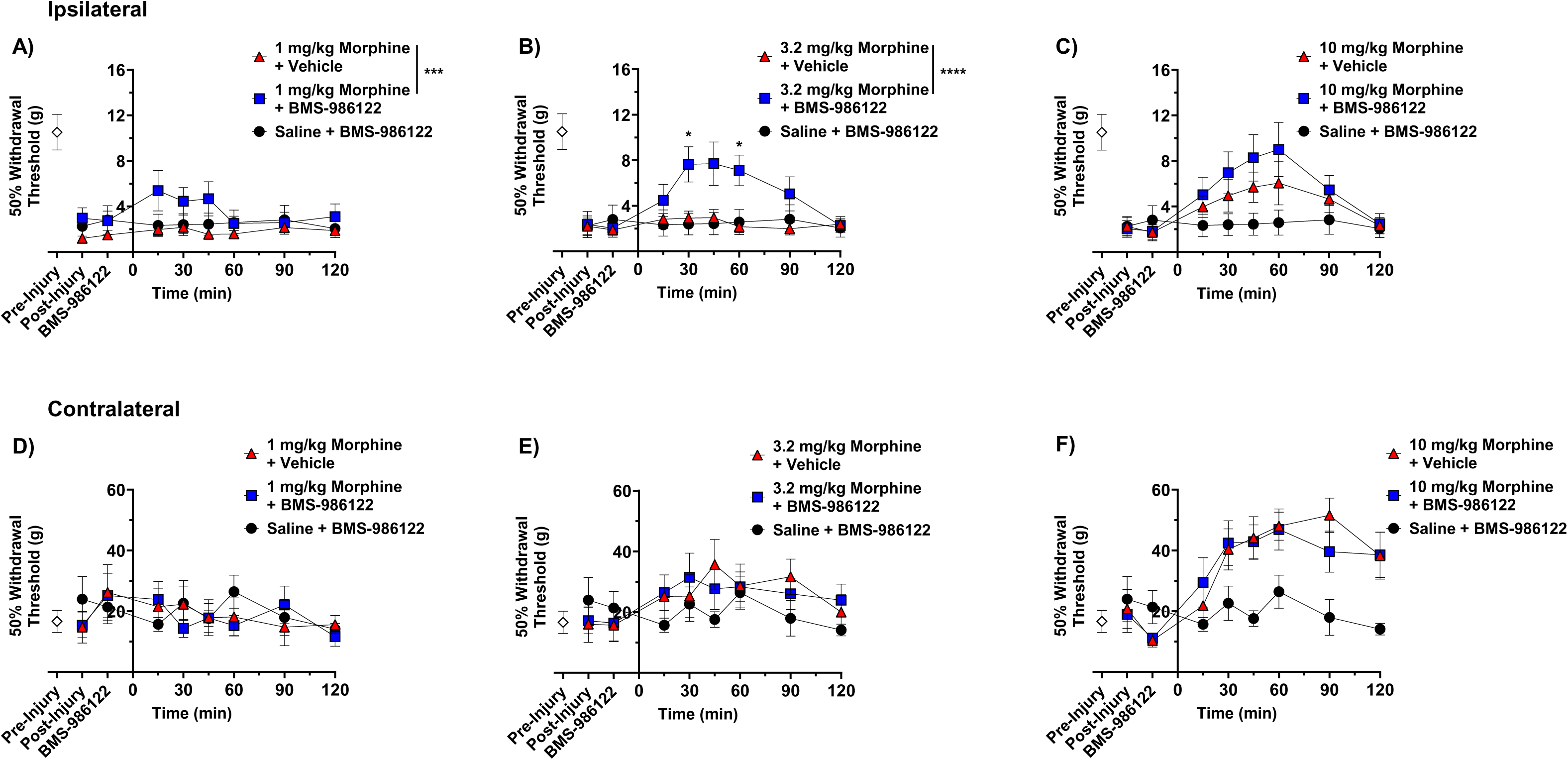
Effect of BMS-986122 on morphine-induced reduction in tactile hypersensitivity after nerve injury. Rats with tactile hypersensitivity in the injured hind paw after SNI received BMS-986122 (10 mg/kg, s.c.) as a 15-minute pretreatment to subcutaneous morphine at a dose of 1 mg/kg (**A**), 3.2 mg/kg (**B**), or 10 mg/kg (**C**). 50% Withdrawal thresholds were determined using von Frey filaments and the up-down method prior to injury (Pre-Injury) and prior to BMS-986122 administration (Post-Injury), 15 minutes after BMS-986122 administration (BMS-986122), and for 2 hours after morphine administration. Thresholds were simultaneously measured in the contralateral, uninjured hind paw after mice received 1 mg/kg (**D**), 3.2 mg/kg (**E**), or 10 mg/kg (**F**) morphine. Groups were compared using a repeat-measures two-way ANOVA with Tukey’s post hoc test, with significance reported between morphine in the presence of vehicle and morphine in the presence of BMS-986122 at each time point (n=8, 4 females and 4 males). * p < 0.05, *** p < 0.001, **** p < 0.0001). Data are presented as mean ± S.E.M.

Due to the risks associated with full opioid agonists, the use of the partial agonist buprenorphine is becoming preferable in the clinic.^26^ As PAMs have been shown to enhance the analgesic efficacy of partial agonists *in vitro* and *in vivo*,^7,9,11,33^ we hypothesized that BMS-986122 would increase the maximal response to buprenorphine in the SNI model. Buprenorphine (0.01 mg/kg) did not change the tactile threshold, either alone or in the presence of BMS-986122 (**Fig. 3A**). However, the action of a low efficacious dose of buprenorphine (0.032 mg/kg) was enhanced by BMS-986122 (**Fig. 3B**). A higher dose of 0.1 mg/kg of buprenorphine did not further improve the effect of the opioid alone, as it reached a ceiling in line with its partial agonist activity (**Fig. 3C**). On the other hand, in the presence of BMS-986122, this dose of buprenorphine further reversed the hypersensitivity (**Fig. 3C**), showing an enhancement in the efficacy of the opioid. Similarly to (R)-methadone and morphine, buprenorphine increased thresholds on the uninjured paw, with no difference in the presence or absence of BMS-986122, except for the 90 min time point in animals treated with 0.032 mg/kg buprenorphine (**Fig. 3D-F**).

**Figure 3.**
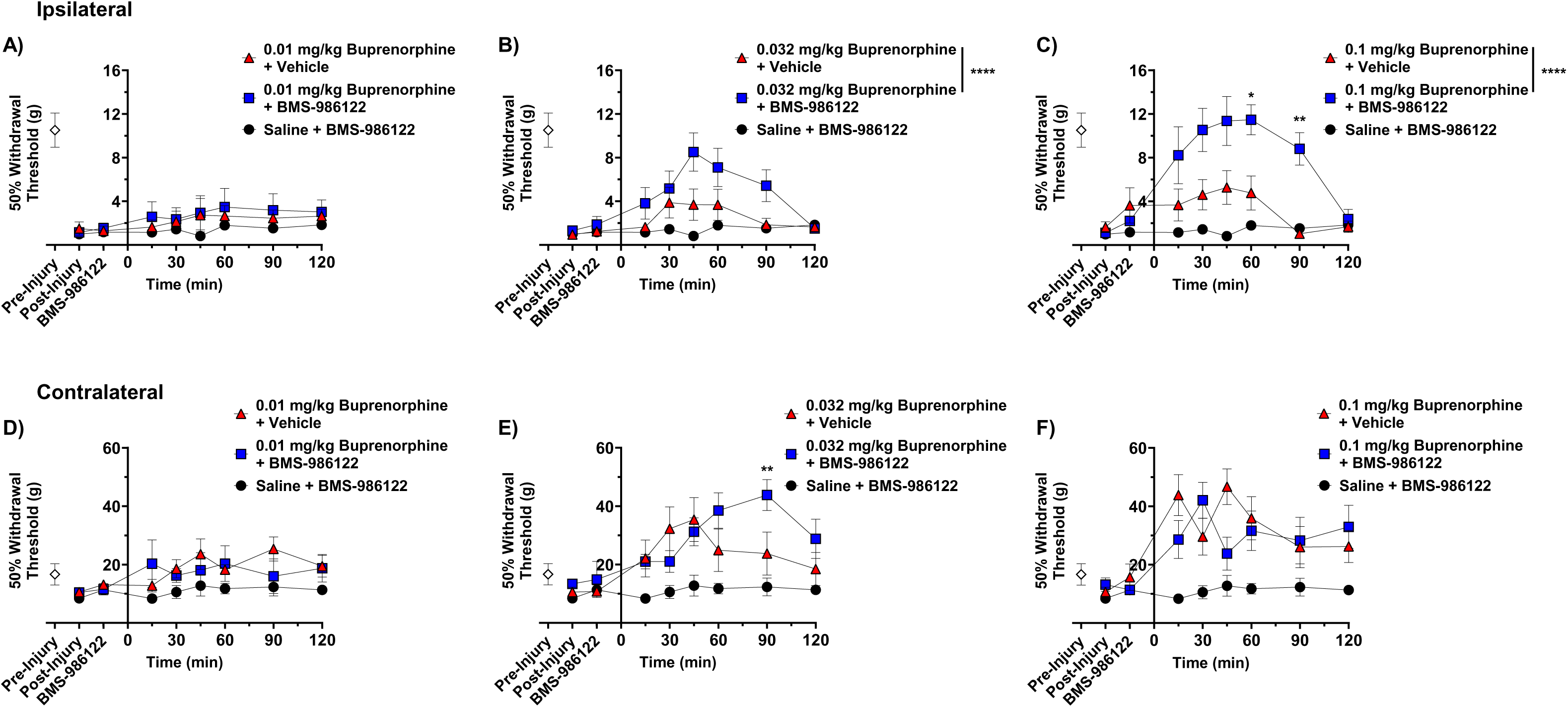
Effect of BMS-986122 on buprenorphine-induced reduction in tactile hypersensitivity after nerve injury. Rats with tactile hypersensitivity in the injured hind paw after SNI received BMS-986122 (10 mg/kg, s.c.) as a 15-minute pretreatment to buprenorphine at a dose of 0.01 mg/kg (**A**), 0.032 mg/kg (**B**), or 0.1 mg/kg (**C**). 50% Withdrawal thresholds were determined using von Frey filaments and the up-down method prior to injury (Pre-Injury) and prior to BMS-986122 administration (Post-Injury), 15 minutes after BMS-986122 administration (BMS-986122), and for 2 hours after buprenorphine administration. Thresholds were simultaneously measured in the contralateral, uninjured hind paw after mice received 0.01 mg/kg (**D**), 0.032 mg/kg (**E**), or 0.1 mg/kg (**F**) buprenorphine. Groups were compared using a repeat-measures two-way ANOVA with Tukey’s post hoc test, with significance reported between buprenorphine in the presence of vehicle and buprenorphine in the presence of BMS-986122 s at each time point (n=8, 4 females and 4 males). * p < 0.05, ** p < 0.01, **** p < 0.0001. Data are presented as mean ± S.E.M.

Opioids are not typical first-line treatments for chronic neuropathic pain.^5^ Therefore, we compared the effects of low dose opioid treatments with or without BMS-986122 to gabapentin, a standard of care medication. Administration of 30 mg/kg gabapentin to rats with SNI did not significantly increase 50% withdrawal thresholds compared to vehicle-treated animals, but 100 mg/kg was fully efficacious in reversing the hypersensitivity (**Fig. 4A**). Thus, the degree of effectiveness opioids in the presence of BMS-986122 was similar to that seen with gabapentin. (**Fig. 1C, 2C, 3C**). Gabapentin, as with opioids, also increased contralateral thresholds at the highest dose tested (**Fig. 4B**).

**Figure 4.**
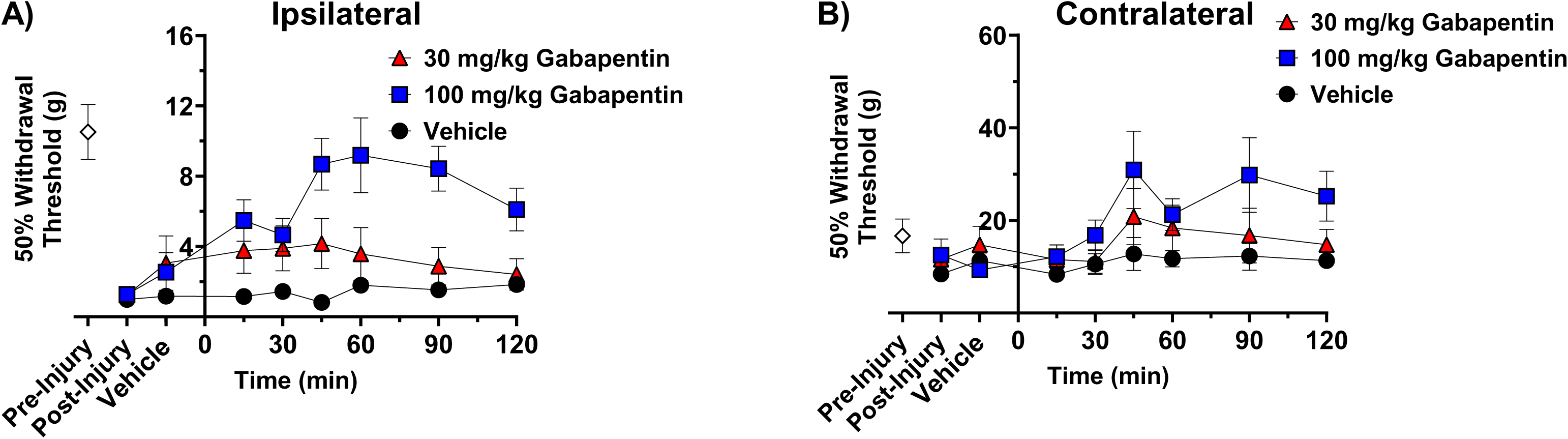
Effect of gabapentin on tactile hypersensitivity induced by nerve injury. Rats with tactile hypersensitivity in the injured hind paw after SNI received gabapentin at a dose of 30 or 100 mg/kg s.c. (**A**). 50% Withdrawal thresholds were determined using von Frey filaments and the up-down method prior to injury (Pre-Injury), post-injury, and for 2 hours after gabapentin administration. Thresholds were simultaneously measured in the contralateral, uninjured hind paw (**B**). Data are presented as mean ± S.E.M (n=8, 4 females and 4 males).

### Neuroma Pain

While SNI is a well-established rodent model of chronic neuropathic pain, deep mononeuropathies to the sciatic nerve are rare.^34^ Therefore, we tested the efficacy of BMS-986122 in a different model of neuropathic pain, namely tibial neuroma, which provides an alternative model of traumatic injury.^28^ The neuroma was formed by transection of the left tibial nerve by the knee, transposition to the dermis and suturing to secure the nerve ending.^28^ This affords a neuroma that is highly sensitive to tactile palpation. The number of palpations with a 26g von Frey filament until a reflexive response occurs was used as a measure of hypersensitivity, defined as the “neuroma score”. This closely resembles the Tinel sign, a distinct response related to neuroma pain in patients.^16^ The number of palpations necessary to elicit a response from a human neuroma is recorded as a Tinel score, with 1 being highly sensitive and 6 as insensitive. After injury and neuroma formation in the rat model, a neuroma score of 1-2 was recorded at the neuroma site, representing high sensitivity to tactile stimuli (**Fig. 5**). As in the SNI model, (R)-methadone was administered s.c. and a dose of 10mg/kg BMS-986122 s.c. was given as a pretreatment.

**Figure 5.**
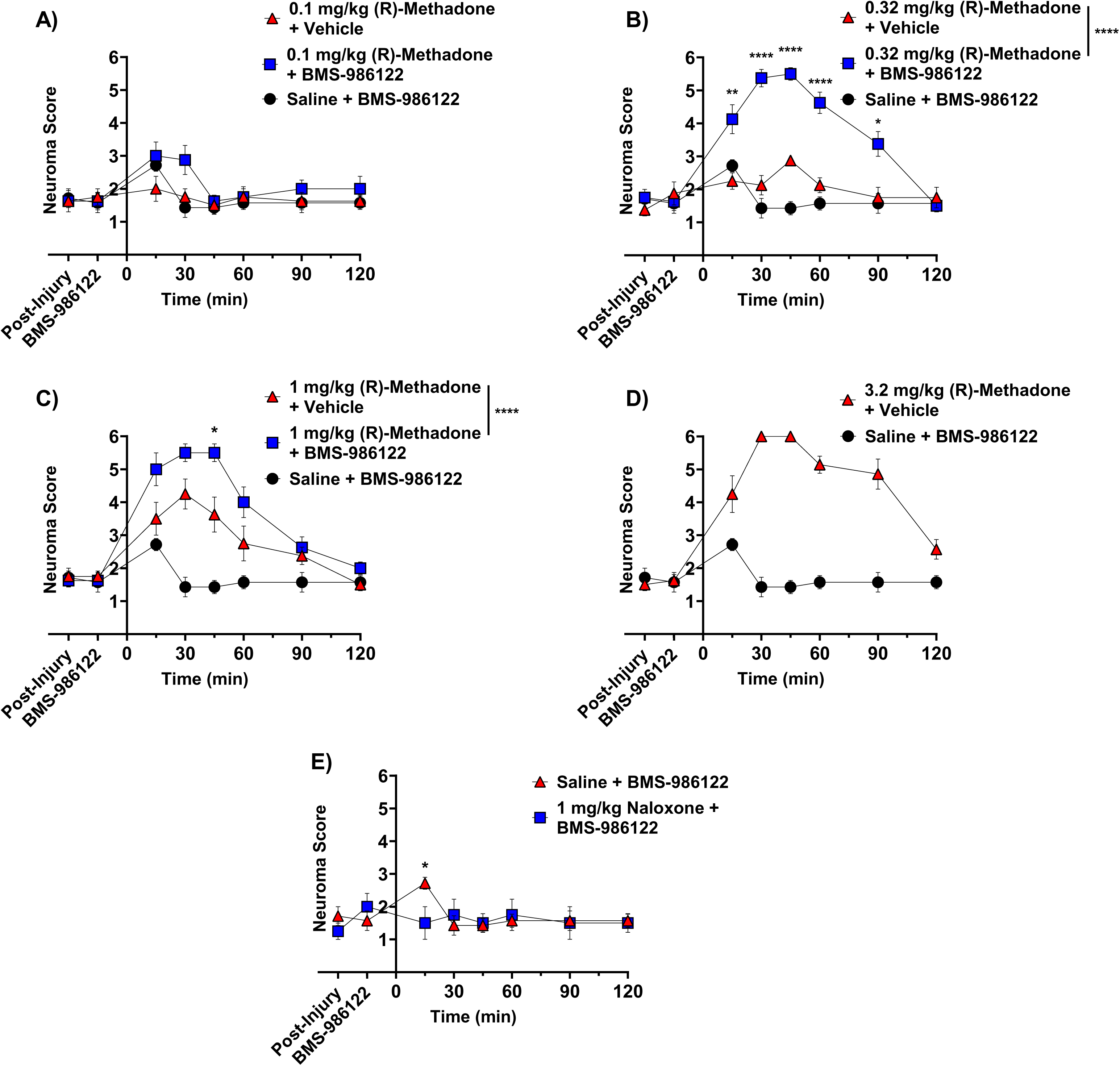
Time-course of effects of BMS-986122 on inhibition of neuroma hypersensitivity by (R)-methadone. Rats with tibial neuroma respond to tactile stimulation of the neuroma, as measured by number of stimuli necessary to evoke a reflexive response (Neuroma score). Rats received BMS-986122 (10 mg/kg) as a 15-minute pretreatment to subcutaneous (R)-methadone at a dose of 0.1 mg/kg (**A**), 0.32 mg/kg (**B**), or 1 mg/kg (**C**). Animals received 3.2 mg/kg (R)-methadone (with BMS-986122 vehicle) to determine maximal opioid response (**D)** or BMS-986122 alone (E). The neuroma score in each group was determined prior to BMS-986122 administration (Post-Injury), 15 minutes after BMS-986122 administration (BMS-986122), and for 2 hours after (R)-methadone 2 hours 15 min after PAM, administration. Groups were compared using repeat-measures two-way ANOVA with Tukey’s post hoc test, with significance reported between methadone in the presence of vehicle and methadone in the presence of BMS-986122 (n=8, 4 females and 4 males). To test for the opioid component effects of BMS-986122 shown in (E), 1 mg/kg naloxone (s.c.) was administered 15 mins prior. Groups were compared using multiple Student’s t-tests (n=4, 2 males and 2 females). * p < 0.05, ** p < 0.01, *** p < 0.001, **** p < 0.0001 (n = 8). Data are presented as mean ± S.E.M.

Administration of low dose (R)-methadone (0.1 mg/kg or 0.32 mg.kg) did not increase the neuroma score, representing no reversal of hypersensitivity, but animals did show a significant response in the presence of BMS-986122, reaching maximal effect at 0.32 mg/kg (R)-methadone when animals did not respond after 6 palpations. (**Fig. 5A, 4B**). Administration of 1 mg/kg (R)-methadone alone did increase the neuroma score and reached a maximal response in the presence of BMS-986122 (**Fig. 5C**). Methadone alone reached a maximal response at 3.2 mg/kg (**Fig. 5D**). Since 0.32 mg/kg (R)-methadone was fully effective in the presence of BMS-986122 this suggests the modulator improves the potency of (R)-methadone by approximately 10-fold. Animals given 3.2 mg/kg (R)-methadone alone showed respiratory distress and had to be rescued with naloxone. Notably, no respiratory problems were noted in animals given lower doses of (R)-methadone (0.032 – 1.0 mg/kg) in the presence of BMS-986122, even though maximal antihypersensitivity was observed.

In the tibial neuroma model, administration of BMS-986122 in the absence of (R)-methadone slightly and transiently increased the neuroma score 30 minutes after administration. This effect was blocked by administration of 1 mg/kg naloxone (**Fig. 5E**).

## Discussion

Current drug treatments for chronic neuropathic pain are inadequate for patients, either due to difficulty in phenotypic profiling, heterogenous diagnostic criteria, and/or modest efficacy of the therapies themselves.^5^ The lower efficacy of opioids in chronic neuropathic pain is thought to be a result of reduction in MOR expression and general neuroplasticity after injury. In rodent models of nerve injury, MOR expression decreases in the periphery^31,35,36^ and in the central nervous system,^37–39^ particularly within the descending pain modulatory systems.^40^ Similar changes in receptor expression also occur in humans with both peripheral and central nerve damage.^41^ Here, we report that pretreatment with BMS-986122, a PAM acting at MOR, enhances the activity of low doses of the MOR agonists in rat models of both mononeuropathy and neuroma pain. The combination of low-dose opioid with BMS-986122 afforded responses comparable to the current standard of care, gabapentin. We hypothesize the ability of BMS-986122 to enhance the action of opioids in the chronic nerve injury models is likely by enhancing their activity at MOR^10^ to compensate for the reduced levels of MOR expression. Moreover, given that we have previously shown the PAM does not promote the action of opioids acting at MOR to cause constipation, respiratory depression or reward,^11^ this suggests MOR-PAMs may be effective as opioid-sparing agents in the management of neuropathic pain.

In rats with SNI, the ipsilateral hind paw develops high sensitivity to pressure as measured with von Frey filaments. Low doses of (R)-methadone or morphine were largely ineffective in reversing this hypersensitivity, but combined with BMS-986122, these low doses of opioids fully reversed the hypersensitivity. The hypersensitivity of the rat neuroma to touch with von Frey filaments was reversed by high doses of (R)-methadone (3.2 mg/kg), but these doses caused respiratory distress in the animals. However, in the presence of BMS-986122 the potency of (R)-methadone was increased 10-fold, in agreement with our *in vitro* findings^27^, such that 0.32 mg/kg fully reversed the hypersensitivity of the neuroma. In contrast, this potency enhancement did not apply to the effects on respiration as none of the animals treated with methadone plus BMS-986122 showed evidence of respiratory depression. Given that *in vitro*, in overexpressed cell systems, we observed a similar enhancement of MOR activity, the reason for this selective effect on nociceptive responses but not respiration is not immediately clear. Nonetheless, the results do indicate that the combination of opioids and PAM might show an increased safety margin and adds evidence to previous data that BMS-986122 promotes the antinociceptive actions of opioids but not their respiratory depressant actions.^11^

Partial agonists are safer than more efficacious agonists such as morphine or methadone as they are much less likely to lead to abuse or cause life threatening respiratory depression. As such, clinical opioid therapy has begun to embrace the use of the partial agonist buprenorphine to treat pain while limiting opioid abuse and overdose.^42–44^ Buprenorphine administered alone only afforded a small degree of reversal of the hypersensitivity response in rats with SNI and reached a ceiling effect at 0.1 mg/kg, in agreement with its partial agonist activity. In contrast, in the presence of BMS-986122, an increase in the maximal response to buprenorphine was seen such that the opioid now fully reversed the tactile sensitivity. This is supported by our *in vitro* finding that PAMs increase the maximal response to partial agonists.^11^ Most buprenorphine prescriptions are for chronic low back pain.^43,44^ Should our findings translate to humans, then opioid PAMs could prove a valuable source of clinical adjuvants to increase the efficacy of buprenorphine in humans without enhancing negative side effects, providing wider application of buprenorphine to chronic pain.

In the tibial neuroma model, a small beneficial effect of BMS-986122 administered alone was observed. This action was blocked by naloxone, confirming opioid receptor involvement. BMS-986122 has been shown to act as an “ago-PAM”, i.e. a PAM that itself has agonist activity. However, this is dependent on a high level of MOR expression, as seen in cell systems,^33^ so is not a probable reason for its activity, as *in vivo* expression is far lower. On the other hand, BMS-986122 has been shown to increase MOR receptor activation by opioid peptides *in vitro*^10^ and to enhance the antinociceptive effects of endogenous opioids *in vivo* in acute models of pain.^7^ Based on these findings, it is likely that the PAM is increasing the activity of endogenous opioids at the orthosteric site on MOR to an extent that an antihyperalgesic effect can be observed. It is known that endogenous opioid peptide levels change in certain chronic pain conditions in patients^45^ and in rats with inflammation,^46^ and may possibly increase in rats with neuroma as well. However, studies in animals suggest that opioid peptide expression does not change or may even decrease over longer periods following nerve injury and neuroma formation, both in the periphery and the spinal cord.^47,48^ On the other hand, it is possible that the changes in peptide expression occur in higher centers following neuroma formation that could explain the minor effect of BMS-986122 alone. In contrast, opioid peptide expression is reportedly not increased following SNI,^49,50^ which may explain why BMS-986122 alone shows no activity in this model.

This study, as a preliminary assessment of the efficacy of opioid PAMs in neuropathic pain, contains several limitations. A reflexive response to tactile stimulation was the only behavioral outcome used, yet this only represents a part of the phenotype of a chronic pain state. While tactile hypersensitivity is common across neuropathic pain disorders,^39^ further study of additional non-reflexive pain behaviors could provide additional insight into the extent to which MOR PAMs enhance the activity of opioids in chronic pain states. For example, previous work with BMS-986122 in mice showed enhancement of nest building after an acute noxious stimulus to the abdomen, supporting the effectiveness of the compound in non-evoked pain.^7^ As opioids will induce sedation in rats, it is possible the reflexive responses measured may be confounded by sedation. The PAM did not increase opioid effects in the contralateral paw, though, suggesting that the effects are not due to general sedation. In addition, while males and females were compared to test for sex differences, the sample sizes were only appropriate for testing overt effects. Smaller sex effects may still contribute to these results. Finally, although BMS-986122 has been shown not to enhance respiratory depression, constipation, or reward in mice,^11^ these outcomes have not been tested in rats and were not examined in this study. Conversely, the lack of observable respiratory distress in animals given low-dose (R)-methadone plus BMS-986122 suggest the respiratory depressant actions of (R)-methadone are not promoted. However, due to solubility problems all our experiments were performed using 10 mg/kg BMS-986122 and we were unable to examine higher doses which could have uncovered unwanted side effects.

In conclusion, the current data demonstrate that MOR PAMs could be a potential treatment strategy for opioid sparing in the management of chronic pain states, particularly in resistant neuropathic pain states like neuroma pain. Moreover, these modulators could increase the efficacy of the safer partial agonists in the management of chronic neuropathic pain. The capacity of PAMs, like BMS-986122, to widen the therapeutic window of opioid medications, and so reduce the opioid burden on patients with chronic neuropathic pain, would have a positive clinical benefit while limiting risks to this patient population.

## Prior Presentations

Clements, B.M., Kerrigan, K.E., Hsuing-Lee, C., Berberoglu, I., Kemp, S.W.P., Traynor, J.R. Positive Allosteric Modulator BMS-122 Enhances the Anti-Allodynic Action of Opioid and Non-Opioid Analgesics in Rats with Nerve Injury. Poster presentation delivered at the International Narcotics Research Conference. Bologna, Italy July 2025.

Clements, B.M., Kerrigan, K.E., Kemp, S.W.P., Traynor, J.R. Opioid Positive Allosteric Modulators Enhance Methadone-Mediated Anti-Allodynia in a Rat Model of Peripheral Nerve Injury. Poster presentation delivered at the Society for Neuroscience Annual Meeting. Washington, D.C. November 2024.

Clements, B.M., Kerrigan, K., Kemp, S.W.P., Traynor, J.R. Enhancement of Opioid Analgesia by Positive Allosteric Modulation of the μ-Opioid Receptor in a Rat Model of Chronic Neuropathic Pain. Poster presentation delivered at the American Society for Pharmacology and Experimental Therapeutics. Arlington, VA. May 2024.

## Summary Statement

A positive allosteric modulator of the mu-opioid receptor enhances the anti-allodynic effects of clinically used opioids in neuropathic pain.

## Acknowledgements

We would like to thank Dr. Mark Bicket and Dr. Steven Harte for their guidance and support during the execution of these studies and the preparation of this manuscript.

## Funding

National Institutes of Health grant R37-DA33397 and R42-(JRT)

National Institutes of Health and Michigan Institute for Clinical and Health Research grant T32-TR004764 (BMC)

Department of Defense grant CDMRP W81XWH-21-1-0771 (SWPK)

## Competing Interests

JRT holds a NIH STTR grant with Eleven Therapeutics Corp. BMC and SWPK declare that they have no competing interests.

## Data and materials availability

All data are available in the main text or the supplementary materials. Original data files are available upon request.

## Abbreviations and Acronyms

MOR: µ-opioid receptor
PAM: positive allosteric modulator
s.c.: subcutaneous

## Notes

### Competing Interest Statement

The authors have declared no competing interest.

